# The cholinergic pesticide imidacloprid impairs contrast and direction sensitivity in motion detecting neurons of an insect pollinator

**DOI:** 10.1101/295576

**Authors:** Elisa Rigosi, David C. O’Carroll

## Abstract

Cholinergic pesticides such as the neonicotinoid imidacloprid are the most important insecticides used for plant protection worldwide. In recent decades concerns have been raised about side effects on non-target insect species, including altered foraging behaviour and navigation. Although pollinators rely on visual cues to forage and navigate their environment, the effect of neonicotinoids on visual processing have been largely overlooked. Here we describe a modified electrophysiological setup that allowed recordings of visually evoked responses while perfusing the brain *in vivo*. Long-lasting recordings from wide-field motion sensitive neurons of the hoverfly pollinator, *Eristalis tenax*, revealed that sub-lethal exposure to imidacloprid alters their physiological response to motion stimuli. We observed substantially increased spontaneous firing rate, reduced contrast sensitivity and weaker directional tuning to wide-field moving stimuli. This approach reveals sub-lethal effects of imidacloprid in the visual motion detecting system of an important pollinator with likely implications for flight control, hovering and routing.

## Introduction

Major ongoing debate concerns the off-target effects on animal populations of widely used agrichemicals including neonicotinoid pesticides, such as imidacloprid (recently reviewed in bees in Alkassab and Kirchner, 2017). While targeted at pest species, beneficial arthropods such as pollinating insects are also exposed to these potential threats, both directly (via nectar and pollen of treated plants), and indirectly through exposure at sites of accumulation (e.g. soil, water, nests)(Alkassab and Kirchner, 2017). Neonicotinoids, including imidacloprid (IMI), act agonistically on nicotinic acetylcholine receptors (nAChRs) at synapses in the insect nervous system (Liu *et al.*, 1993; Matsuda *et al.*, 2001). Yet despite being commercially available for more than 20 years, surprisingly little is known about how sub-lethal doses of these chemicals affect the insect nervous system function, particularly in intact individuals.

Prior work on the neurophysiological effects of these chemicals have primarily utilised cultured cells and isolated brain preparations (see for example: Buckingham et al., 1997; Déglise *et al.*, 2002; Barbara *et al.*, 2008; Palmer *et al*., 2013). While these *in vitro* approaches certainly help in identify effective sites of action, they provide little information on how neuronal function is affected in the whole living organism and thus in the presence of relevant external sensory stimuli. Recent attempts to redress this deficiency include a calcium imaging study that recorded odour-evoked responses from the antennal lobe of the intact honey bee and revealed impaired odour processing when the brain was exposed to an acute dose of IMI (Andrione *et al.*, 2016).

The dearth of studies on the effects of IMI on the visual system is surprising given that many parts of the insect visual system are known potential targets for cholinergic agonists. Both cholinergic neurons and nAChRs are widely expressed in the insect optic lobes – both peripherally and centrally (Kreissl and Bicker, 1989; Brotz *et al.*, 2001; Sinakevitch and Strausfeld, 2004; Raghu *et al.*, 2011). Direct evidence for potential effects on visual processing includes acute exposure to sub-lethal doses of imidacloprid, which caused increased cell death in the optic lobes of honey bees (de Almeida Rossi *et al.*, 2013). More recently, visually-evoked responses recorded in a pre-motor neuron, DCMD, from the locust ventral nerve cord showed impaired burst activity in intact animals previously injected with a sub-lethal dose of imidacloprid compared to animals injected with its vehicle (Parkinson *et al.*, 2017). This study is of particular interest as it represents the first and only evidence of impairment *in vivo* in the visual nervous system of an insect due to imidacloprid exposure, although where and how this deficit arises in the upstream optic lobes remains unknown.

Bees fed orally with sub-lethal doses of imidacloprid show alterations of complex visually-guided behaviours such as spatial learning, navigation and homing flights (reviewed in Alkassab and Kirchner, 2017). Ubiquitously, throughout these behaviours, flying insects need to stabilize their head and body position during flight manoeuvres and course control. In dipteran flies, this visually driven flight control is mediated by a well-described class of neurons in the 3^rd^ optic ganglia, the motion-sensitive lobula plate tangential cells (LPTCs). Both *in vitro* pharmacological studies and immunohistochemistry have showed that these class of wide-field direction sensitive neurons in the fly are activated by ACh and express nAChRs in proximity of their dendrites (Brotz *et al.*, 2001; Schmid, 1992; Brotz and Borst, 1996).

Given their well described role during flying manoeuvres and their association with the ACh system, we hypothesized LPTCs as a likely site of action of cholinergic pesticides. To test this, we developed a new preparation that allows us to record extracellularly or even intracellularly from LPTCs in the hoverfly, *Eristalis tenax* while perfusing the brain haemolymph with sub-lethal doses of IMI and during subsequent washout. Our preparation leaves the animal largely intact, allowing us to stimulate the eye with wide-field directional stimuli comprised of moving visual gratings and directly quantify the effects of IMI or its vehicle on electrophysiological responses of LPTCs. We found that when exposed to imidacloprid these cells increased their spontaneous activity, as well as decreased both their contrast sensitivity and directional tuning compared either to their normal state or to animals exposed to a control stimulus the vehicle. Our new preparation thus not only identifies profound effects on the processing of directional motion in a species which is an important pollinator in its own right, it provides a new method for ongoing analysis of other visual pathways or in other species that should valuable to examine possible links between sub-lethal doses of neonicotinoids or other agrichemicals and visual function.

## Materials and Methods

### Experimental Design

We built an *in vivo* perfusion system (Fig. 1) to be able to constantly perfuse and expose the insect brain to a sub-lethal dose of imidacloprid (3.9 μM) and at the same time perform long-lasting electrophysiological recordings from lobula plate tangential cells (LPTCs) of an intact insect pollinator.

**Figure 1:**
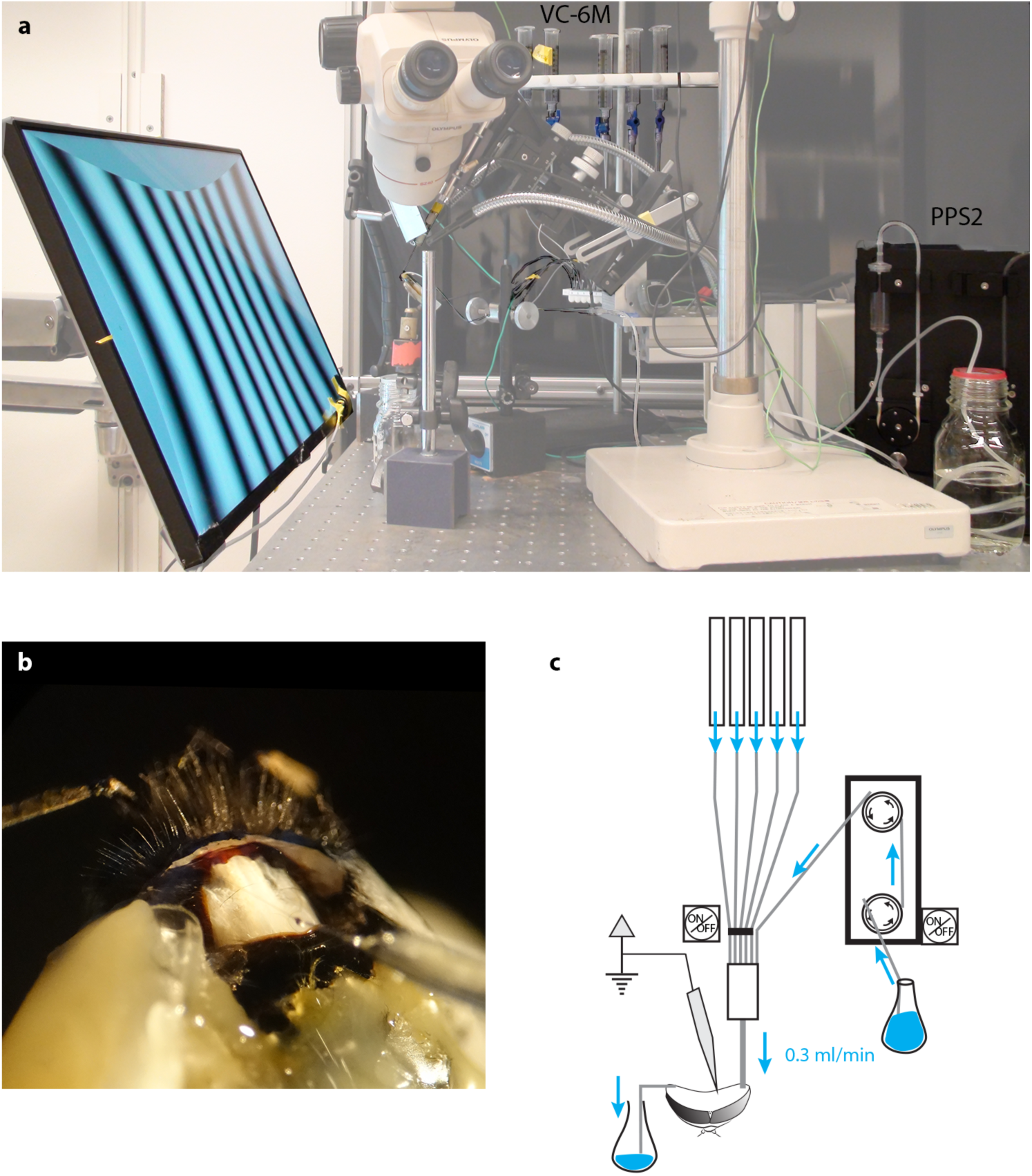
Electrophysiological set up for long lasting recordings in perfused insects. We modified our existing set up to perform electrophysiological recordings while constantly perfuse the insect head capsule. (**A**) A low-noise peristaltic perfusion pump was fed into a micro ML-6 manifold apparatus that received also 5 PE-50 tubing from a 6-valve computer-controlled gravity system, each of this connected to a 20 ml syringe filled with treatment solutions. (**B**) Close up of the insect head showing the inlet tubing, the outlet and the recording electrode (the example shown is in the bumblebee *Bombus terrestris* rather than *Eristalis*). (**C**) Schematic drawing of the perfusion system.

### Animal preparation

Hoverflies [*Eristalis tenax* (Linnaeus, 1758)] were collected outdoors between August and November 2017. From a total of 28 preparations, sufficiently stable recordings for analysis were obtained from two males and eight females. After animal collection on flowers, flies were taken into the lab; if not immediately used, individual flies were kept at 4°C in plastic bags with wet tissue and granulated brown sugar up to 10 days. Flies maintained in the fridge for longer than 2 days were brought to room temperature for ~1h every 48h to allow them to feed and clean themselves and the wet tissue with sugar replaced. Immediately before the experiment the insect was fed *ad libitum* with granulated brown sugar on a wet tissue.

Flies were inserted into a pipette tip, cut at its narrow end to approximate the diameter of the head, and the thorax and mouthparts fixed with low-temperature melting compound of 1:1 wax:rosin, tilting the insect head about 45° forward to optimize neural recording from the lobula plate tangential cells. First the reference electrode was inserted at the base of the right eye. Then ~150 mm length of thin polyethylene tubing PE-50 (outside diameter: 0.97 mm, inside diameter: 0.58 mm; Warner instruments, Harvard apparatus) was fixed with wax along the thorax so that the tip was able to perfuse the back of the eye. The wax:rosin compound was used to fill the gaps between the edge of the eye and the thorax in both sides in order to prevent the perfusion liquid to run off the head capsule. Subsequently, the cuticle from the back of the head was gently removed, fat bodies and tracheae removed with tissue paper and forceps. As soon as the brain was exposed the animal was put into the electrophysiological set up, the perfusion tubing connected (see below) and the electrode inserted within 1 minute, to avoid desiccation.

### Perfusion system

A low-noise peristaltic perfusion pump (PPS2, Multi Channel Systems MCS GmbH, flow rate: 0.3 ml/min) was fed into a micro ML-6 manifold (Warner Instruments, Harvard apparatus) that received also 5 PE-50 tubing from a 6-valve computer-controlled gravity system (VC-6M Valve Control System, Warner Instruments, Harvard Apparatus). Each input was connected to a 20 ml syringe filled with treatment solutions (Fig. 1). Each of the channels of the gravity system was height adjusted to match the flow rate of the peristaltic pump (0.3 ml/min). The outlet of the micro manifold was then connected to the animal’s head through the PE-50 tubing as described above.

In our initial experiments with the suction-system provided by the PPS2 pump system to maintain a constant level of fluid in the head capsule, we found vibration and electrical noise to be problematic as the suction system cyclically made and broke contact with the meniscus in the perfused area. Subsequently we solved this problem to maintain a constant fluid level in the head capsule through a capillary-based system whereby we placed the tip of a thin cotton thread on the right side of the head with the other extremity in a glass container to collect the liquid (Fig. 1). The constant capillary action in this thread gently draws fluid away from the perfused area at a rate high enough to allow the entire volume of the interior head capsule to be changed approximately every 10 s (0.3 ml/min). After experimenting with several different threads for this purpose, we found that a washed tea-bag string (Twinings, UK) provided an ideal compromise between size and capillary action.

### Drug delivery

Due to its low solubility in water, a stock solution of imidacloprid (Sigma-Aldrich Sweden AB) was obtained dissolving 1 mg of imidacloprid in 1ml DMSO (Sigma-Aldrich Sweden AB) and maintained at −20°C. On the day of each experiment the stock solution (IMI 3.9 mM) or DMSO alone were then diluted (1:1000) in insect Ringer solution comprising (in mM) the following: NaCl (140), KCl (10), NaH_2_PO_4_ (4), Na_2_HPO_4_ (6), CaCl_2_(H_2_O)_2_ (3), sucrose (90), and adjusted to pH 6.8.

A single test concentration (IMI 3.9 μM) was chosen on the basis of EC50 obtained in previous experiments in isolated insect brains and cultured neurons in honey bees and cockroaches (Buckingham *et al.*, 1997; Déglise *et al.*, 2002; Barbara *et al.*, 2008; Palmer *et al.*, 2013).

### Electrophysiological recordings and visual stimuli

Extracellular recordings were performed at room temperature (23-26°C) using aluminosilicate glass capillaries (SM100F-10, Harvard Apparatus, Holliston, MA, USA) pulled with a P-2000 laser puller (Sutter Instruments, Novato, CA, USA) and filled with 1M KCl solution. The tip of the capillary was gently broken on fine (1000 grit) carborundum paper to obtain a resistance of 1-10 MΩ and subsequently inserted in the lobula complex via a micro-manipulator (Marzhauser-Wetzlar PM-10; step size, 5 μm). The extracellular recordings were digitized at a 10 kHz sample rate, after hardware filtering with low-pass (3 kHz) and high-pass (3 Hz) filters built into the differential preamplifier (BC-03x, NPI, Germany).

The animal sat in front of a high luminance, high speed IPS LCD monitor (Asus ROG Swift PG279Q, 2560 x 1440 pixels; ~380 cd/m^2^; 144 Hz) and wide-field motion-sensitive neurons were identified on the basis of their characteristic responses to wide-field motion in opposite directions (Fig. 2) and weak responses to smaller features. Because of the type of recordings (extracellular) we could only select the direction-sensitive cells on the basis of their spiking activity (see for example in Hausen, 1982) with no regards to graded changes in membrane potentials.

**Figure 2:**
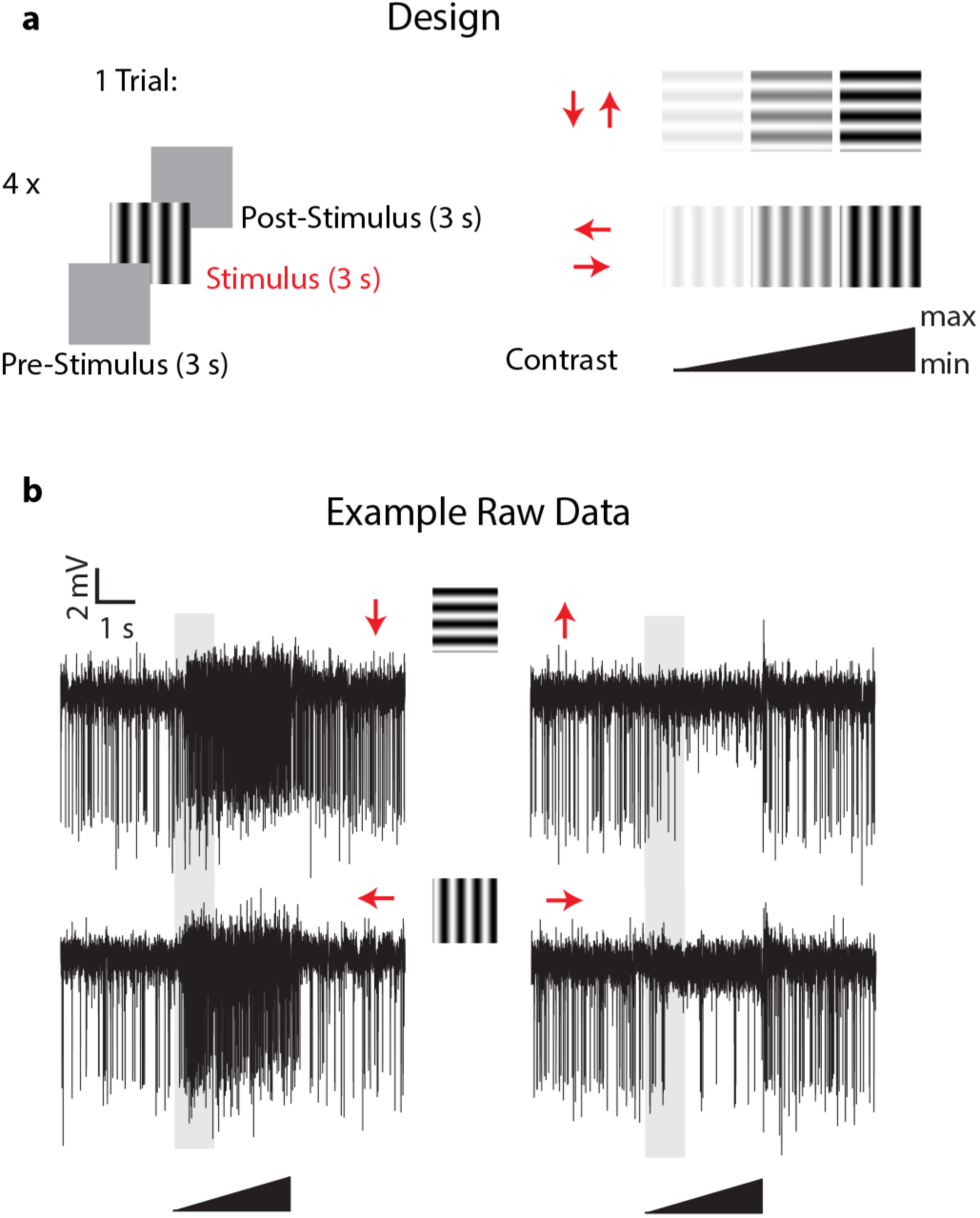
Experimental design and example of LPTCs direction selective responses in *Eristalis tenax* with ongoing perfusion. (**a**) the experimental design consisted of the following visual stimuli: a mean luminance (grey) screen left for 3 s (pre-stimulus period) was followed by a full-screen visual grating stimulus (3 s) in one of the 4 cardinal directions (Temporal frequency: 8 Hz; Spatial frequency: 0.1 cycles/deg) and then the mean luminance screen again for 3 s (post-stimulus response). Four different stimuli with different directions were tested in one trial, and the contrast of each visual stimulus consisted of a linearly increasing ramp of contrast from 0 to a maximum of either 0.5 or 1.0 (see methods for max values). (**b**) Time-course of visual-evoked responses of a trial in a recorded cell when the head of the animal was perfused with Ringer. Red arrows represent the direction of motion of the stimulus used; the 3 s visual stimulus is represented by the black triangle and the shaded area represents the analysed time-window.

Nevertheless the recording electrode tip location, frontal receptive fields, spontaneous rates, responses to motion stimuli and preferred direction (leftwards or downwards) were all consistent with recordings from vertical and horizontal system neurons of the lobula plate. Visual stimuli were presented using custom-written software implemented in Matlab (The MathWorks, Natick, MA, USA) and Psychtoolbox (psychtoolbox.org), with gamma calibration correction. Once the position of the electrode on the brain was identified as suitable for the targeted neurons the perfusion system (with Ringer solution) was turned on so that the head capsule was filled. At the same time the cotton string was carefully added laterally on the right part of the head to suck the solution away and maintain a constant fluid level within the head volume. Once the liquid in the head capsule was rising and in contact with the capillary containing the electrode, the electrical contact was sometimes lost and a good new single unit recording had to be re-established by moving the electrode a few microns.

Once we had established a clear and stable single-unit response from a wide-field motion-sensitive cell with the perfusion system on, the experiment started, with useful recordings then lasting up to 3h using the same recorded unit. Stimuli comprised a mean luminance (blank) screen for 3 seconds (pre-stimulus period), stimulated with a full-screen visual grating stimulus (Temporal frequency: 8 Hz; Spatial frequency: 0.1 cycles/deg) for 3 s and then a return to the mean luminance screen for 3 s (post-stimulus response, Fig. 2). Four directions were tested in 4 different sequential trials (inter-trials interval: 0 s). The direction experiment was replicated about every 7 minutes. The contrast of each grating stimulus was linearly increasing from a Michelson contrast of 0 to 1, except in 3 cells in which the maximum contrast was 0.5 to avoid response saturation due to the high contrast sensitivity of these cells. 30-40 minutes after the experiment started the perfusion system was switched to the treatment (via the multi-valve gravity system; main pump (PPS2) with Ringer set to off, see Fig. 1) and the direction selectivity tested 5 minutes after the perfusion with treatment started. The treatment lasted for a maximum of 1h, after which the main peristaltic pump with Ringer alone was switched back again (multi-valve gravity system off) for 45 minutes to washout the treatment in the head capsule and the direction stimuli tested every 7 minutes through this period.

### Data analysis and statistics

For each animal, raw data were imported into Spike 2 (version 7.02a, Cambridge Electronic Design), digital filtered with a band-pass filter (125.5 Hz, 3k Hz low and high corner frequency, respectively), spike templates created and single spike events identified. Spike templates were identified in the initial recordings for each cell and these same templates were then successfully applied to detect spikes in all subsequent recordings through to the end of the experiment. On the basis of the template shape and their separation using principal component analysis (PCA), for each animal we identified independent single spiking units. We obtained a total of 12 such units from 10 animals. We excluded from the analysis any units that showed an averaged spike frequency below 40 spike/s for low contrast stimuli (Michelson contrast: 0-0.3) in the preferred direction, and thus we used 10 units from 10 different individuals.

For each sample we obtained 2-10 technical replicates for each direction (Fig. 2, 1 Trial) and within each condition (pre-treatment, during treatment and washout). The averaged spike frequency was calculated during pre-stimulus time (spontaneous response, Fig. 3) and during the ramp stimulus (contrast range 0-0.33, see also Fig.2) for each technical replicate and then averaged for each condition (Fig. 3). Effect size for spontaneous response across conditions is reported as Cohen’s *d* coefficient (average ± 95% C.I.).

**Figure 3:**
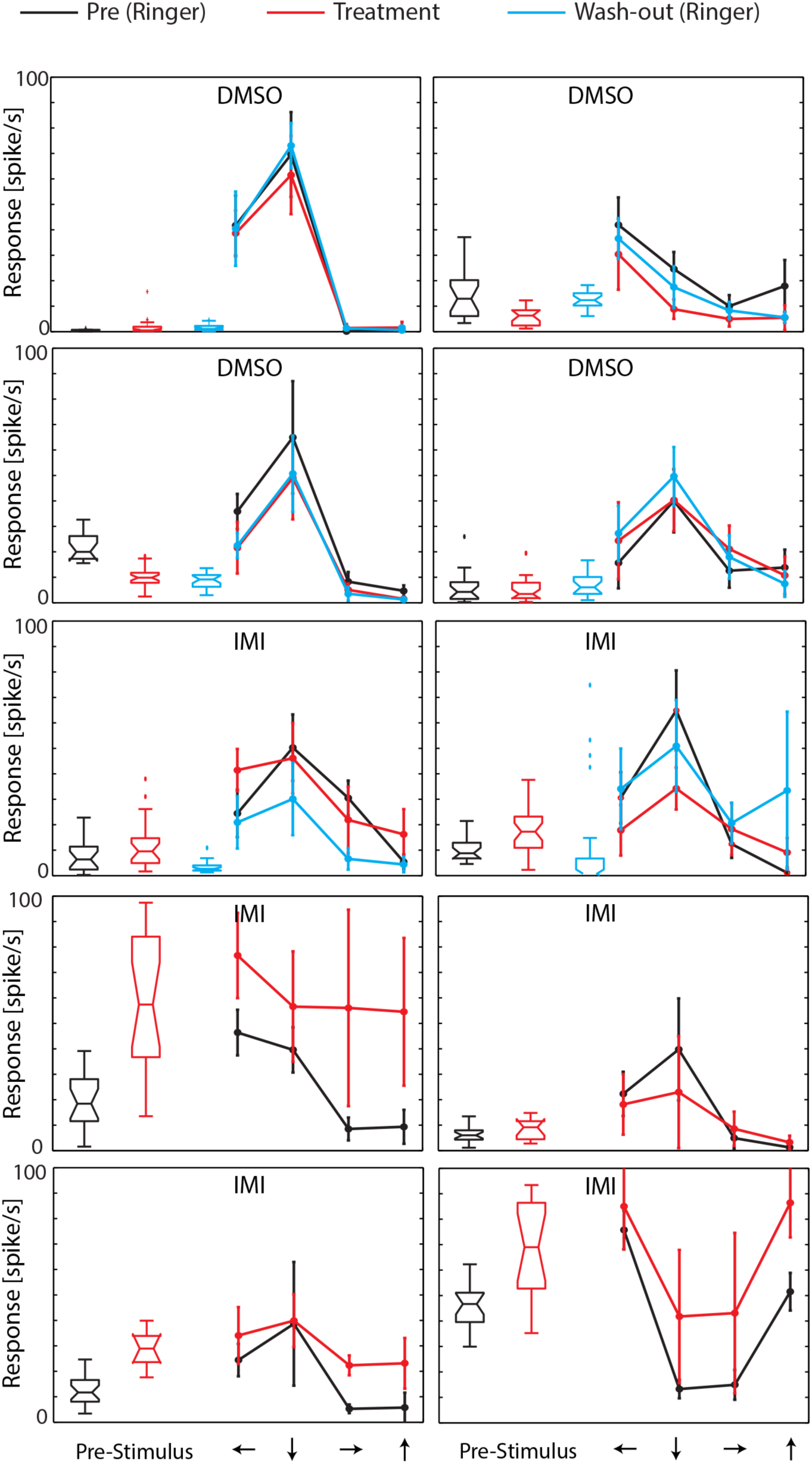
Effects of Imidacloprid (IMI) or its vehicle (dimethyl sulfoxide, DMSO) in 10 lobula plate tangential cells (LPTCs) of the pollinator *E. tenax*. Spontaneous responses (spike/s) for all the recorded units are showed as box-plots. Notched box-plots represent median and 25th-75th percentile, whiskers with maximum 1.5 iqr and outliers are reported. Average response ± SD for the 4 directions in each condition are plotted for each cell (Pre: Ringer, black; during Treatment: either IMI or DMSO, red; washout: Ringer, cyan; *n*=2-10 in each condition).

For each experimental condition, we calculated a directionality index (DI) on the averaged response activity (contrast range 0:0.33) as follows: DI = (P-AP)/(P+AP) with P and AP referring to the averaged activity in the preferred and anti-preferred direction (Fig. 5). Contrast sensitivity was calculated by averaging spiking frequency along the linear contrast ramp in bins of 150 or 320 ms (for stimuli with a maximum contrast of 1 and 0.5, respectively) corresponding to bands of contrast 0.06 wide, in the range between 0 and 0.5 (Fig. 6). For each response in the preferred direction a Weibull function was fit to the data and a simplex search algorithm used to obtain the best-fit values of output range and slope.

**Figure 5:**
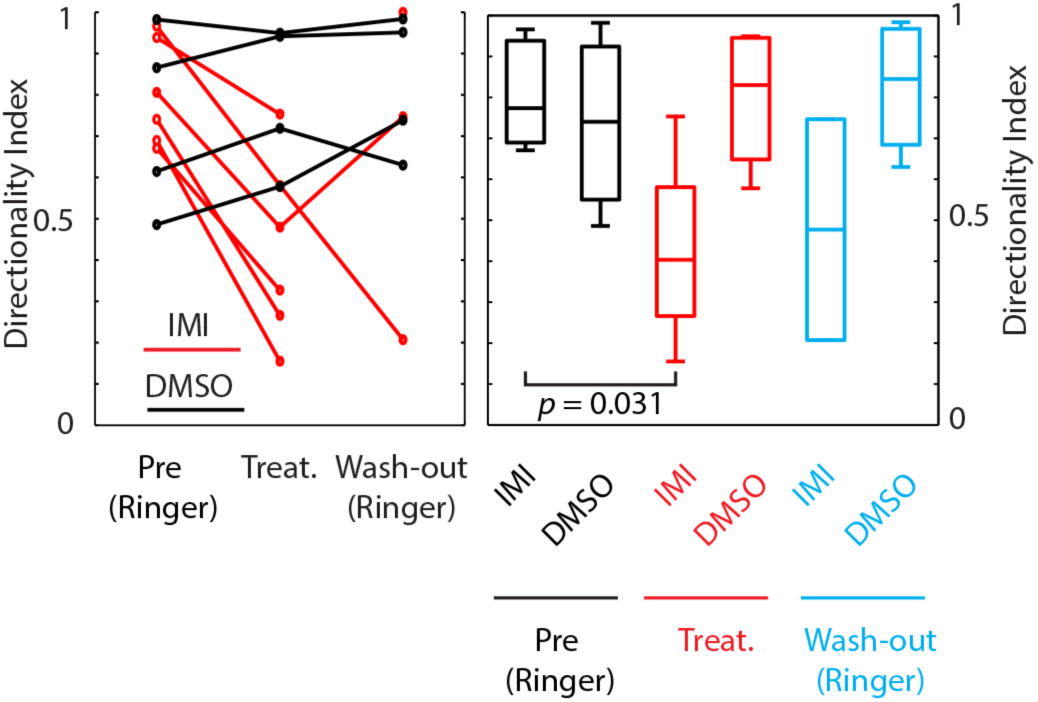
Imidacloprid alters direction selectivity of LPTCs in the fly pollinator *E.tenax* (Left) Directionality index obtained for each cell using the averaged responses in preferred and anti-preferred directions (see methods) in the pre-treatment (Ringer), during treatment and during washout (Ringer) conditions. Cells treated with DMSO (black) are superimposed to ones treated with IMI (red). **(Right)** Box-plots of directionality index for all the conditions of the same data plotted on the left, median, quartile and extremes are plotted.

**Figure 6:**
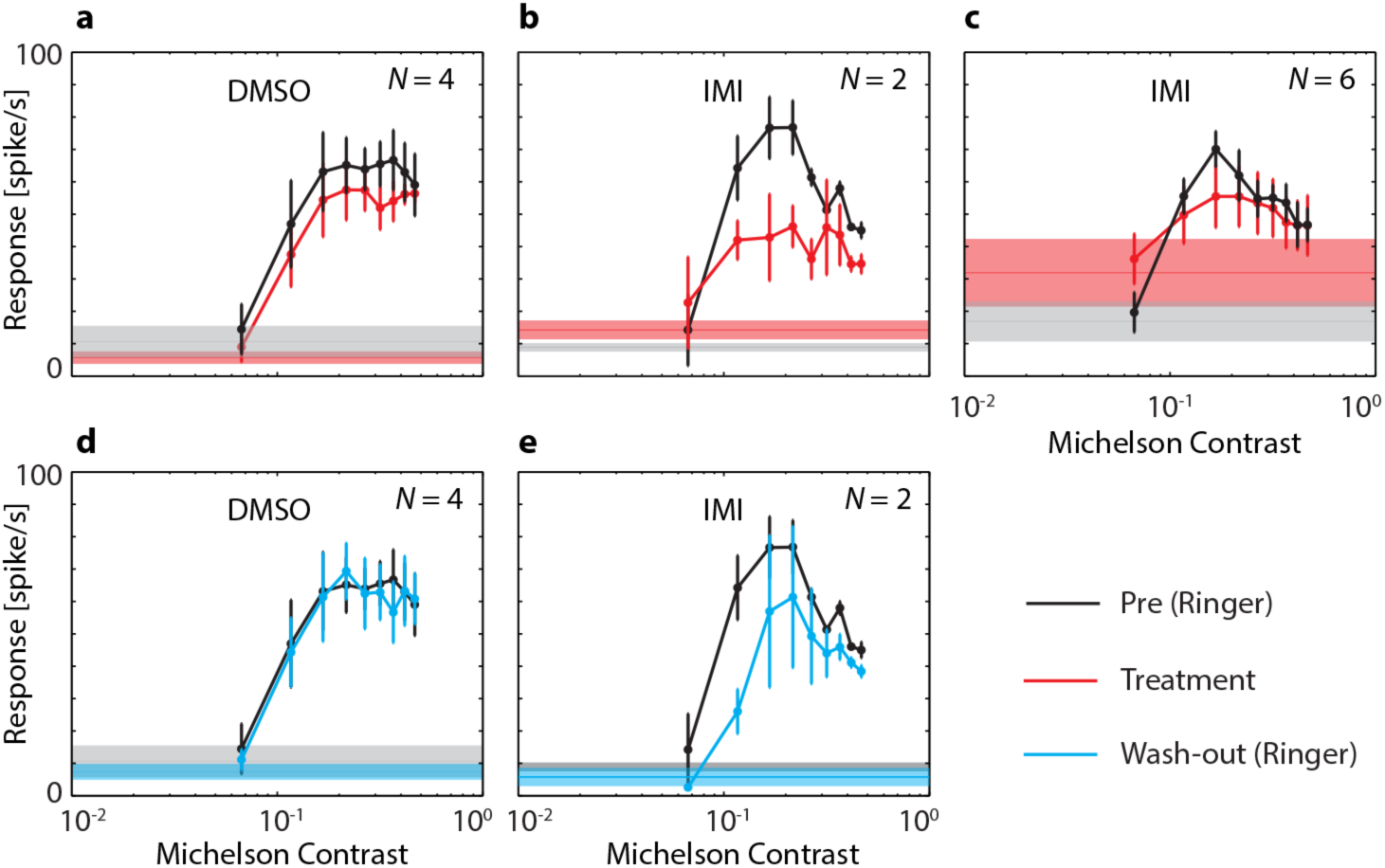
Imidacloprid disrupts contrast sensitivity of LPTCs in *E. tenax*. Average ±SEM responses to different contrasts in cells that are exposed to DMSO (a, d) or to IMI (b,c,e). Contrast sensitivity curve are showed during pre-treatment (perfused with Ringer, black lines) and during treatment exposure (red lines) in the upper part of the figure. C shows pre-treatment and treatment contrast responses in all cells where IMI was used (*N* = 6), while b, e, only show data for experiments where we could also record responses during washout (*N* = 2). To make it more readable in the bottom part of the figure we plotted again the contrast sensitivity in the pre-treatment condition (perfused with Ringer, black lines) this time with the washout (cyan). Spontaneous activity is shown as shaded area (averaged ±SEM) of the same colour of the perfusion condition. Contrast values are mean values of contrast obtained binning approximately 0.06 contrast intervals and responses are averaged absolute spike frequency in the corresponding time windows. Stated *N* values are the biological replicates (individuals) used for each plot.

Due to low number of samples we used non-parametric statistic as follows: Wilcoxon signed Rank Test to compare pre-treatment conditions and Mann–Whitney Rank Sum to compare between treatments (IMI-DMSO), all tests were performed in Matlab.

## Results

### Long-lasting extracellular recordings from the insect lobula in constantly perfused animals

In order to expose the insect brain to sub-lethal concentration of the neonicotinoid imidacloprid we set up a peristaltic pump combined with a computer-controlled multi valve gravity fed system to supply artificial haemolymph to the head capsule of the insect (Fig. 1). This system allows for either intracellular or extracellular recordings from the insect brain while continuously perfusing with a main flow (usually Ringer solution) and up to 5 different treatments to switch into this flow at a relatively high flow rate (0.3 ml/min). The success and duration of the recordings in these conditions depend upon the size and type of neurons targeted for recording, the proximity of the electrode tip to the inlet of the perfusion system and on the type of recordings (intra/extracellular). We used this setup to record from motion-sensitive lobula plate neurons of the hoverfly, *Eristalis tenax*, and investigate how their physiological response to 4 different directional stimuli (Fig. 2) was affected by the exposure to sub-lethal doses of imidacloprid (3.9 μM).

Our application of a low-vibration pump to supply the main saline flow and our novel implementation of a vibration-free capillary suction system permitted brief intracellular recordings from lobula plate neurons (data not shown). However, we found that more stable extracellular recordings from the same class of neurons allowed for longer-lasting recording sessions, up to three hours in healthy cells. This allowed us to obtain visual-evoked responses during a resting state (pre-treatment), during the treatment and during subsequent washout of the treated agent, all within the same unit. However, even with our low-vibration delivery and suction system, this type of recording remains highly challenging and the success rate (long-lasting, healthy recording with good signal-to-noise ratio, see for example Fig. 2) was about 1 out of 3 preparations. Of these healthy recordings we successfully treated 6 animals with imidacloprid and 4 with its vehicle, dimethyl sulfoxide (DMSO). While the healthy recordings with the vehicle were able to last through the subsequent washout (~ 3 h) in all cases, we lost good electrophysiological contact after treatment in 4 out of 6 animals treated with IMI.

### Imidacloprid increased spontaneous responses of LPTCs

We first asked whether the spontaneous response of the neurons was affected by treatment exposure. In the pre-treatment condition, the two groups (IMI, DMSO) showed no significant difference in spontaneous spike rates (Fig. 3 and 4, *p*=0.609, Mann–Whitney Rank Sum Test). However, following exposure either to IMI or to its vehicle, DMSO, these neurons showed significantly different spontaneous activity (*p* = 0.019, Mann–Whitney Rank Sum Test) but in opposite directions: IMI increased the spontaneous firing rate (with a large effect size, Cohen’s d ± 95% C.I.: −1.03 ± 0.4), while the vehicle, DMSO, slightly decreased it (Cohen’s d: 0.55 ± 0.9). During washout the cells exposed to DMSO returned to similar levels of spontaneous activity as in pre-treatment (Cohen’s d: 0.20 ± 1.1), while in the 2 cells exposed to IMI the treatment was reversible and went slightly below the pre-treatment firing rate (Cohen’s d: 0.50 ± 0.8; Fig. 3 and 4).

**Figure 4:**
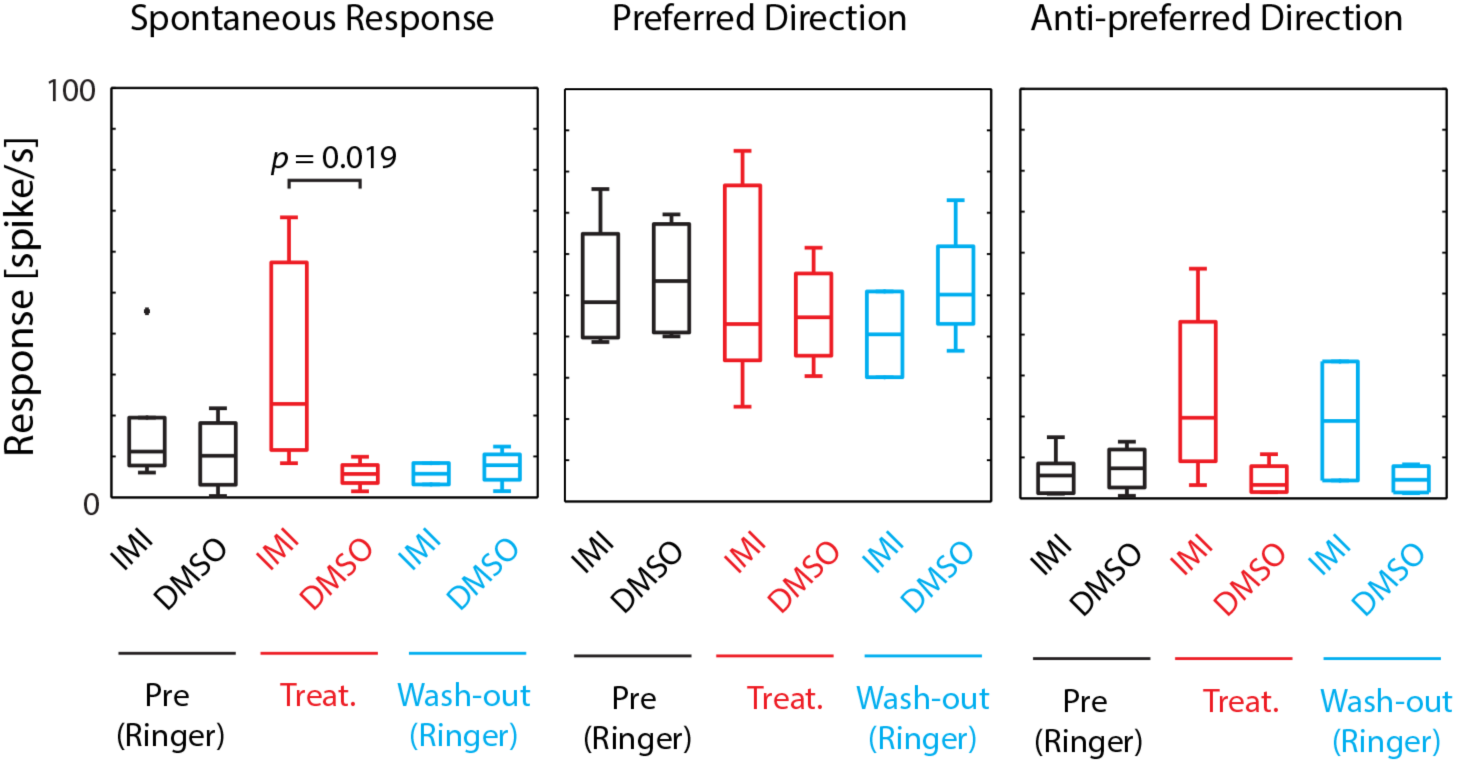
Imidacloprid increases LPTCs spontaneous responses in the fly pollinator *E.tenax.* Box-plots of averaged absolute response during pre stimulus time (spontaneous response, left panel) during a visual grating stimulus moved in the preferred direction (middle) and anti-preferred direction (right panel) for all treatment and conditions (DMSO: *N* = 4; IMI: *N* = 6 in pre and treatment conditions, *N* = 2 in washout). Box-plots represent median and 25th-75th percentile, whiskers with maximum 1.5 iqr and outliers are reported.

### Imidacloprid altered directional tuning and contrast sensitivity

An inclusion criterion for our classification of the recorded neurons as LPTCs was that they were initially motion opponent i.e. their spontaneous activity was inhibited by stimuli in the anti-preferred direction (Fig. 2). Interestingly, we observed a general increase in the response to anti-preferred direction stimuli when cells were exposed to IMI, sometimes linked to a decreased response to motion in the preferred direction (Fig. 3 and 4) suggesting that the general direction sensitivity of the neuron might have been affected. To investigate this further, we calculated a directionality index in each cell for every condition (Fig. 5), whereby a value of 1.0 indicates complete inhibition by anti-preferred directions, and a value of 0 would indicate no difference between the preferred and anti-preferred responses. When pooled together we observed a decreased direction selectivity in cells treated with IMI (Fig. 5; *p* = 0.031, Wilcoxon signed Rank Test) but not in those treated with DMSO (Fig. 5, *p* = 0.250, Wilcoxon signed Rank Test). In the latter treatment, in fact, the directionality index was stable throughout the 3 conditions of the experiments (black dots in Fig. 5, left panel).

We finally asked whether IMI affects the contrast sensitivity of LPTCs. Each stimulus presented consisted of a linear ramp of increasing contrast (Fig. 2), so responses that commence earlier within the presentation period indicate higher sensitivity to the stimulated pattern (O’Carroll *et al.*, 1997). This allowed us to analyse the contrast sensitivity of the response in the preferred direction by temporally binning the responses into small contrast ranges (Fig. 6). Similar contrast sensitivities were observed in the pre-treatment conditions for the two groups of neurons, with a clear departure from spontaneous response levels at a threshold contrast of ~0.05 (i.e. contrast sensitivity > 20) and saturation of the response at contrasts above a contrast of 0.2. At higher contrasts still, the response declines due to motion adaptation to the extended stimulus. During IMI treatment, contrast sensitivity curves changed dramatically, starting from a much higher spontaneous level. While the contrast required to reach response saturation was similar, the overall response level at high contrasts was clearly reduced. Hence while contrast sensitivity did not differ in the pre-treatment conditions (*p* = 0.3524, Mann–Whitney Rank Sum Test, Fig.6 a,c) the IMI treated response shows a profound reduction in output range (response gain) rather than an obvious change in threshold (contrast gain), compared with those treated with DMSO (Fig. 6 a–c, *p* = 0.009, Mann–Whitney Rank Sum Test). While DMSO caused a slight decrement in saturation level, this was consistent with reduced spontaneous activity and was completely reversed during washout (fig 6 a,d). In the 2 cells where we were able to complete washout following IMI-treatment (fig. 6 b,e) the cells still showed a substantially reduced output range, despite the spontaneous activity returning to low levels, indicating that contrast sensitivity was only partially restored during the washout.

## Discussion

We used electrophysiological recordings combined with a peristaltic perfusion system to investigate the effects of sub-lethal doses of the widely used neonicotinoid, imidacloprid, on the wide-field motion sensitive cells of the pollinator fly *Eristalis tenax*. The fly lobula plate tangential cells (LPTCs) are a well-known model for the study of motion detection (reviewed in Silies *et al.*, 2014). These neurons normally show increased activity when a wide-field moving stimulus is presented in a specific direction and inhibited when the motion is presented in the opposite direction (see Fig. 2). When 3.9 μM imidacloprid was perfused in the head capsule of the animal, the recorded cells showed significantly elevated spontaneous rates but weaker responses to normally-excitatory or inhibitory stimuli. i.e. both their directional selectivity and contrast sensitivity were altered, indicating a generally reduced ability to encode attributes of stimuli to which these cells are normally sensitive.

LPTCs get their inputs from direction-selective columnar neurons that process locally retinotopic motion signals from the periphery (Silies *et al.*, 2014; Douglass and Strausfeld, 1996). As has been theoretically described, direction opponent responses such as observed in Fig. 2 arise from the computation derived from dendrites of LPTCs and their inputs where both inhibitory GABA and excitatory nACh receptors are jointly present (Brotz *et al.*, 2001; Sinakevitch and Strausfeld, 2004). Indeed both GABA and nACh receptors are seen in close proximity of LPTCs (Brotz *et al.*, 2001) and are functionally activated by ACh and its agonists (Schmid, 1992; Brotz and Borst, 1996). Thus it seems plausible that in addition to any indirect impairment in other cholinergic neurons downstream from the LPTCs, the cholinergic pesticide imidacloprid might act directly on the LPTCs to cause their increased firing rate in the absence of any visual stimulus (increased spontaneous rate compared to cells treated with DMSO). This could be either through direct action on nAChRs or through the opening of voltage-gated Calcium channels (see for example Jepson *et al.*, 2006).

When the latter were blocked in cultured cholinergic cells of *D. melanogaster* larvae, in fact, the imidcacloprid-induced activation of these cells was significantly reduced (Jepson *et al.*, 2006).

The increase in spontaneous firing rates we observed is consistent with previous analysis in isolated brains or cultured cells, where bath application or local injection of imidacloprid caused cell depolarization and increased inward (cation) currents (Buckingham *et al.*, 1997; Déglise *et al.*, 2002; Barbara *et al.*, 2008; Palmer *et al.*, 2013). An increase in spontaneous spike frequency in LPTCs was also shown when nicotine was applied in preparations where the animals were intact and responding to a wide-field motion stimulus (Schmid, 1992). Interestingly, Schmid found that nicotine increased spontaneous spike rates in the motion sensitive H1 cells of the fly *Calliphora erytrocephala* but did not affect direction selectivity (Schmid, 1992). This is, to some extent, similar to what we observed in our experiments: when wide-field visual stimuli were presented the cells continued responding to the salient stimuli direction they were tuned to, but with a decreased response strength and with an increased response in the anti-preferred direction (i.e. decreased motion opponency). Imidacloprid, in fact, did not abolish the stimulus-evoked responses in our case as seen previously in projection neurons (Andrione *et al.*, 2016) and in the visual pre-motor neuron in locusta (Parkinson *et al.*, 2017). Why then we do not observe a shut down of the responses? This might be explained by a dose-dependent effect, if, for example, our concentration triggers increased spontaneous responses that did not completely saturate the activity of the neuron. Alternatively, it could reflect a different affinity of the nACh receptor in the LPTC pathway for imidacloprid (Dupuis *et al.*, 2011) compared to the one in the above mentioned studies. However, other parts of the visual system might also indirectly affect the response we record in LPTCs; lamina, medulla and lobula neurons do express nAChR (Kreissl and Bicker, 1989; Brotz *et al.*, 2001; Raghu *et al.*, 2011) as well as a centrifugal modulation from lamina to photoreceptors that has been hypothesized to be glutamatergic or cholinergic (Zheng *et al.*, 2006).

Our results reveal an impairment in direction sensitivity in cells treated with imidacloprid showing both lower strength in the preferred direction (Fig. 5) and by increased response to the anti preferred direction. The latter might be caused by a general increase of the spontaneous response driven by the excitatory ACh current at the LPTC inputs and/or by a reduced inhibition possibly due to a deactivation of GABA-induced currents that are imidacloprid-dependent, as observed in Kenyon cells (Déglise *et al.*, 2002). We also found that IMI impaired the contrast sensitivity of LPTCs, in particular reducing the range over which variations in the stimulus strength produce modulation of the response. This might be partially explained by a general decrement in response activity (i.e. what we observe in the preferred direction) and/or by an impaired contrast sensitivity in the downstream processes, or by other indirect mechanisms, for example through activation of octopaminergic neurons that do have a role in changing contrast output range of the animal (Jung *et al.*, 2011). Either way, the reduced output range we observed indicate a profound effect on the ability of these neurons to encode relevant attributes of the input stimulus, whether it be the direction or the speed of a moving pattern.

Importantly, IMI is not readily soluble in water and needs to be dissolved in a suitable vehicle for delivery either to the physiological preparation we used here, or in field application of this chemical to plants as a systemic insecticide. The vehicle that we used, DMSO, also seems to affect LPTC responses (see spontaneous rate Fig. 3, 5 and contrast sensitivity, Fig. 6a). This was particularly evident in the contrast sensitivity curves where a slight decrement in contrast range is subsequently restored during washout (Fig. 6d). As far as we are aware, effects of DMSO have not been described in behavioural or neurophysiological experiments when testing pesticides. However decreased neural activity due to DMSO has been reported: high concentrations of DMSO, in fact, causes silencing of responses in sensory neurons in *Locusta migratoria*, for example (Theophilidis and Kravari, 1994), as well as induces changes in the permeability of neuron membranes (example in the visual system: Weckström and Laughlin, 2010).

An obvious question that arises from this kind of experiment is whether the dose we used is field-relevant. We used a concentration comparable with imidacloprid EC50 values previously described in insect neurons. These values change on the basis of the preparation and type of neurons, but are in a range between 0.3-23 μM at least as described in the honey bee and in *Periplaneta* cholinergic neurons (Buckingham *et al.*, 1997; Déglise *et al.*, 2002; Barbara *et al.*, 2008; Palmer *et al.*, 2013). A limitation for us in selecting a field-relevant dose is the lack of studies assessing the toxicity of imidacloprid through this specific route of exposure, i.e. applied directly within the head capsule. However, Moffat and colleagues (2015) showed that imidacloprid reaches similar concentration in the brain to the ingested source, within 3 days of chronic exposure (Moffat *et al.*, 2015). A clear picture of what a relevant field dose is nevertheless missing, with imidaclopird concentration in nectar and pollen found on site probably 2 orders of magnitude lower than the concentration we used (Blacquière *et al.*, 2012). However, a similar concentration to the one we used was found, for example, in pollen stored in apiaries in 2010 (Mullin *et al.*, 2010) and concentrations up to 2 orders of magnitudes higher than we used were measured in guttation drops of seed treated plants (Girolami *et al.*, 2009). Nonetheless, these values might be an underestimate of what the brain of the insect is exposed to considering multiple sources of exposure and accumulation. Furthermore this picture is complicated by the observation that the secondary metabolites during chronic exposure showed vastly higher toxicity than acute exposure to imidacloprid itself, no doubt contributing to the lethality of these chemicals in field exposure (Suchail *et al.*, 2001). We did not use such metabolites in our preparation but they would be expected to act on the same receptors (Palmer *et al.*, 2013; Nauen *et al.*, 2001).

Finally, we should consider what effect the changes we observe might have on the animal behaviour. While IMI likely has many other effects on other neuronal pathways than we observed in LPTCs, we can at least speculate as to possible behavioural impairments on the basis of what has been described for the fly motion detection system and LPTC outputs. These neurons are involved in visually driven behaviours intrinsically linked to the detection of motion in specific local directions, e.g. information related to optic-flow, the perceived movement of the world driven by ego-motion. From LPTCs, direct connections project to motor neurons involved in neck and head movements, fundamental for flight-stabilization and gaze control (Gronenberg *et al.*, 1995); as well as indirect projections involved in the looming-detection pathway important for avoidance behaviours (de Vries and Clandinin, 2012). Moreover, motion-information sent through the posterior ventrolateral protocerebrum is presumably used by higher-centres to accomplish behaviours such as landing, flight-speed regulation as well as ‘visual odometry’ for measuring distances (Srinivasan, 1999). Studies of the LPTCs in *Eristalis* indeed reveal that non-linear adaptation in these neurons provides an excellent basis for encoding velocity of optic flow in natural scenes (Barnett et al., 2010) and thus potentially contributing to the kind of visual odometry that plays a key role in navigation to and from the food source in a variety of insect species.

Moreover, motion information encoded in LPTCs has also been shown to be relevant for the detection of objects against a moving background, such as the one the insect experiences while flying through the environment (Fenk *et al.*, 2014; Mertes et al., 2014). Using a behavioural assay in *Drosophila melanogaster*, for example, LPTCs inputs (T4-T5 cells) have been shown to be necessary for object-fixation and discrimination against a high-gain moving background (Fenk *et al.*, 2014). Even more convincingly, Mertes and colleagues (Mertes et al., 2014) recorded from motion-sensitive cells in the bumblebee while presenting visual stimuli reconstructed from their learning flights in an experimental arena. The visual evoked responses from these neurons indeed revealed their role in the detection of landmarks presented in the arena (Mertes *et al.*, 2014). Interestingly, in the past, imidacloprid-induced impairments in honey bee navigation system have been reported: bees fed with sub-lethal doses of IMI showed longer foraging trips as well as incorrect homing flights (Schneider *et al.*, 2012; Fischer *et al.*, 2014), and our findings might at least partially contribute towards the neurophysiological mechanisms involved in such impairments.

To summarize, our data revealed a disrupted direction selectivity and contrast sensitivity in LPTCs after direct IMI application to the brain and a mechanism by which cholinergic pesticides might act in the brain of flying pollinators to effect motion-guided behaviours. Ultimately, our experimental setup will allow future investigation to identify where and how the visual system is affected by neonicotinoids under conditions in which the animal still perceive and respond to environmental relevant sensory stimuli.

## Acknowledgments

We thank the Wenner-Gren Foundation, the Royal Physiographic Society of Lund and the Swedish Research Council (Vetenskapsrådet VR 2014-4904) for their valuable support.

## Author contributions

E.R. and D.C.O. designed the study, analysed data and contributed to the manuscript writing; E.R. performed the experiments.

## Competing interest

The Authors declare no competing interests

